# Withaferin A reduces pulmonary eosinophilia and IL-25 production in a mouse model of allergic airways disease

**DOI:** 10.1101/2024.08.26.609636

**Authors:** Kevin Agner, Victoria L. McQuade, Jack Womble, Marissa A. Guttenberg, Sanat Phatak, Jennifer L. Ingram

**Author notes:** Corresponding author contact information: Sanat Phatak, MD, DM, MRCP, Diabetes Unit, KEM Hospital Research Centre, KEM Hospital, Pune, India.

## Abstract

Several studies report that ashwagandha, a traditional Ayurvedic supplement, has anti-inflammatory properties. Type 2 (T2) asthma is characterized by eosinophilic airway inflammation. We hypothesized that allergen-induced eosinophilic airway inflammation in mice would be reduced following administration of Withaferin A (WFA), the primary active phytochemical in Ashwagandha. C57BL/6J mice were given 10 total intra-peritoneal injections of 2 mg/kg WFA or vehicle control, concurrent with 6 total intranasal administrations of 50 µg house dust mite extract (HDM) or saline control over 2 weeks. We observed that treatment with WFA reduced allergen-induced peribronchial inflammation and airway eosinophil counts compared to mice treated with controls. In addition, we observed that treatment with WFA reduced lung levels of interleukin-25 (IL-25) but increased lung gene expression levels of its co-receptor, *Il17ra*, in HDM-challenged mice compared to HDM-challenged mice that received the vehicle control. This study pinpoints a potential mechanism by which WFA modulates allergen-induced airway eosinophilia via the IL-25 signaling pathway. Future studies will investigate the effects of WFA administration on lung eosinophilia and IL-25 signaling in the context of chronic allergen-challenge.

## Introduction

*Withania somnifera*, commonly known as Ashwagandha, is a woody shrub from the family Solanaceae (Della Porta et al. 2023). It is traditionally used in Ayurvedic medicine for numerous ailments, and most notably has been reported to have neuroprotective, anticancer, and anti-inflammatory properties (Jung et al. 2022; Majeed et al. 2023; Afonso et al. 2023). The foremost active ingredient of Ashwagandha is withaferin A (WFA) (Jung et al. 2022; Patel et al. 2019).

Asthma is a heterogenous chronic disease characterized by airway hyperresponsiveness (AHR), inflammation, and remodeling (Moore et al. 2010). Allergic asthma features Type 2 (T2) cytokine-mediated airway inflammation and chronic airway eosinophilia (Oppenheimer et al. 2022). Mice exposed to acute or chronic allergen challenge exhibit elevated eosinophil levels in bronchoalveolar lavage fluid (BALF) and increased peribronchial eosinophilic inflammation compared to control animals (Briode and Sriramarao 2001; Cates at al. 2004). Eosinophils are activated in the airway by T2 cytokines released from allergen-injured airway epithelium, including interleukin-33 (IL-33) (Cayrol and Girard 2018) and interleukin-25 (IL-25), which signals through a receptor comprised of the interleukin-17 receptor A (IL-17RA) and B (IL-17RB) subunits (Nejman-Gryz et al. 2021).

We hypothesized that allergen-induced lung eosinophilia and release of pro-inflammatory cytokines in the airway would be reduced following administration of WFA in a model of acute allergic airways disease in mice.

## Materials and methods

Animal experiments were approved by the Duke University Institutional Animal Care and Use Committee under protocol #A041-20-02 (approved 03/06/2020).

Six-week-old male and female C57BL/6J mice (n = 19, Jackson Laboratories) were treated with intranasal (i.n.) house dust mite (HDM) extract (Greer Laboratories, #XPB91D3A2.5, Lot 394845) at a dose of 1.25 µg/µL (50 µg/dose), or equivalent volume phosphate-buffered saline (PBS) 3 days/week for 2 weeks. Concurrently, a subset of the mice (n = 3-4 per group) received intraperitoneal (i.p.) injections of 2 mg/kg WFA (Millipore-Sigma, #681535), or equivalent volume of the vehicle (VC; 5% tween-80, 2% dimethylsulfoxide [DMSO] in 1x PBS) 5 days/week for 2 weeks. On days with both i.p. and i.n. doses, mice received the i.p. dose 30 min prior to receiving the i.n. dose (see schematic Figure 1A).

**Figure 1.**
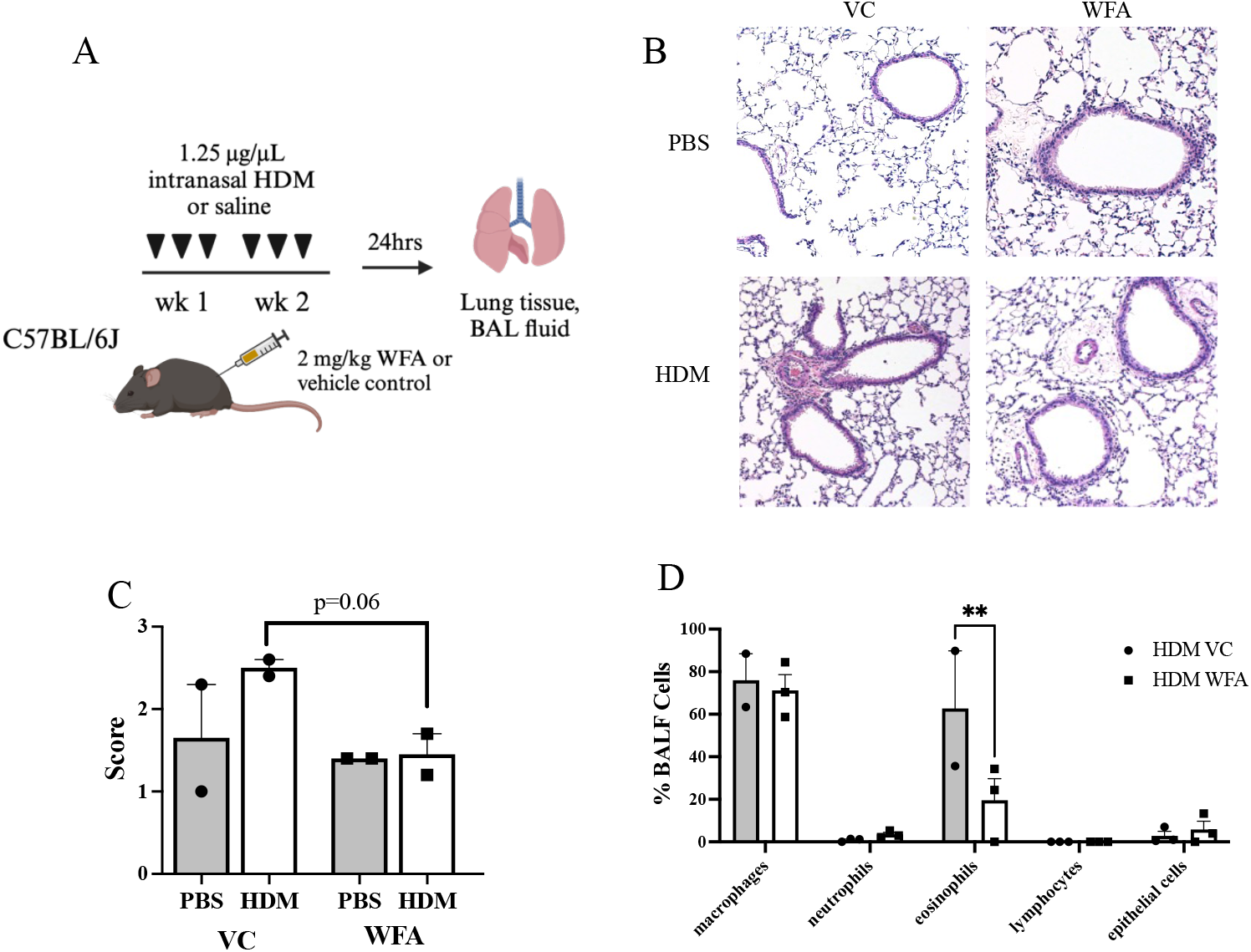
Airway eosinophilic inflammation is reduced with WFA in HDM-challenged mice. (A) Schematic diagram of the mouse model of acute intranasal house dust mite allergen (HDM)-induced allergic airways disease combined with intraperitoneal Withaferin A (WFA) or vehicle control (VC) injection. Control mice were challenged with intranasal phosphate buffered saline (PBS). Bronchoalveolar lavage fluid (BALF) and lung tissue harvest occurred 24 h following the last administrations of HDM/PBS and WFA/VC. Created with BioRender.com. (B) Representative images of H&E-stained lung sections from all 4 treatment groups. (C) Mean H&E scores ± SEM depicting depth of peribronchial staining for n=2 mice per treatment group. (D) BALF differential cell counts in HDM-challenged mice treated with VC or WFA (n = 2-4 mice per group). Data are presented as mean percentage of total BALF cells ± SEM. **p<0.005 for eosinophils in BALF of both treatment groups in HDM-challenged mice.

Mice were sacrificed 24 h after last i.n. dose via lethal injection with 1.5 mg/kg urethane (Millipore-Sigma #U2500). Sacrifice was confirmed via blood draw. BALF was collected via instillation and withdrawal of 1 mL PBS 3 times. The right lung was tied at the bronchus and the medial, lower, and accessory lobes were snap-frozen in liquid nitrogen, while the upper lobe was placed into 1 mL TRI-reagent (Millipore-Sigma, #T9424). The left lung was inflated with 10% formalin and tied off at the bronchus, and the bronchial tree, heart, and left lung were excised in one piece and placed into 5 mL 10% formalin.

Fixed lungs were excised from the bronchus after 24 h and embedded in paraffin. N = 2 lungs per group were sectioned at 5 µm and mounted to slides for staining with hematoxylin and eosin (H&E). Histological scoring was performed as described in Ihrie et al. (2022).

Snap-frozen lobes were macerated in 1 mL PBS using a Tissue-Tearor immersion blender (Biospec Products, Inc., model 985370-395), for 3 s at 20,000 rpm repeated 3 times. The blender was washed between samples by running at 25,000 rpm for 10 s in 70% ethanol and then deionized water before blotting with a paper towel. Samples were centrifuged at 10,000 rpm for 10 min at 4°C to remove cell debris. Lobes in TRI-reagent were similarly macerated and total ribonucleic acid (RNA) extraction proceeded according to the manufacturer’s protocol.

BALF was centrifuged at 1,200 rpm for 10 min at 4°C. The supernatant was removed, and the cell pellet was resuspended in 500 µL red blood cell lysis buffer (Invitrogen, #00430054) and incubated on ice for 5 min. Samples were centrifuged again at the same speed, the buffer was removed, and the pellet was resuspended in 500 µL PBS. Cells were counted via Cellometer K2 (Nexcelom Bioscience) and 25,000 cells/sample were mounted on slides using a Cytospin 4 (ThermoFisher Scientific) at 800 rpm for 3 min. Slides air-dried overnight and then were stained with EasyIII stain (Azer Scientific, ES904, ES905, ES906). Differential cell counts were determined by counting macrophages, eosinophils, lymphocytes, neutrophils, and epithelial cells on cytospin slides imaged at 10× magnification. 200-250 cells were scored per slide.

Whole lung protein concentration was determined via bicinchoninic acid (BCA) assay (Pierce, #23225), and samples were normalized to 15 µg total protein. IL-33 and IL-25 concentration was determined via Enzyme-Linked Immunosorbant Assay (ELISA) kit according to manufacturer’s protocol (IL-33: R&D Systems, DY362605; IL-25: Invitrogen, 88-7002-22).

Whole lung ribonucleic acid (RNA) concentration was determined via NanoDrop One (ThermoFisher Scientific). 50 ng/µL complementary DNA (cDNA) per sample was prepared with a high-capacity reverse transcriptase cDNA kit (Applied Biosystems, #4368814). 50 ng cDNA/sample was assayed using a QuantStudio6 Flex (Applied Biosystems) using TaqMan primers specific for murine *Il17ra* (Mm00434214_m1) and *Gapdh* (Mm99999915_g1) (ThermoFisher Scientific). Fold change in lung *Il17ra* expression in HDM-challenged mice was calculated with the delta Ct method normalizing to PBS as the control treatment and *Gapdh* expression.

Statistical analyses were performed in GraphPad Prism 9 or JMP Pro 17 (SAS, Cary, NC). Outliers were tested with the Robust Regression and Outlier Removal (ROUT) method (Motulsky and Brown 2006) and removed where appropriate. The mouse allergen challenge groups were compared using parametric or non-parametric tests accordingly (one or two-way ANOVA, Kruskal-Wallis, or *t*-test), with appropriate post-test, to evaluate significance. P<0.050 was considered significant.

## Results

Increased inflammation was observed in HDM-challenged mouse lung sections as compared to PBS-challenged mouse lung sections from the VC group (Figure 1B, C left). However, HDM-challenged mice treated with WFA exhibited a reduction in peribronchial inflammation compared to HDM-challenged mice treated with vehicle (VC) (Figure 1B, C right). No difference was observed in PBS-challenged mice treated with WFA compared to PBS-challenged mice treated with VC (Figure 1B, C). In addition, BALF eosinophil counts are reduced in HDM challenged, WFA-treated mice compared to HDM-challenged, VC-treated mice (Figure 1D). PBS-challenged mice exhibited only macrophages in BALF and demonstrated no differences between VC and WFA treatment groups (data not shown).

Lung tissue levels of IL-33 were increased in all HDM-challenged mice compared to PBS-challenged mice, with no difference observed between WFA-and VC-treated mice (Figure 2A). However, lung tissue levels of IL-25 were reduced in HDM-challenged, WFA-treated mice compared to HDM-challenged, VC-treated mice (Figure 2B). A reduction in lung IL-25 levels in HDM-challenged, WFA-treated mice compared to PBS-challenged, WFA-treated mice was also observed (Figure 2C). In addition, *Il17ra* expression was increased in HDM-challenged, WFA-treated mice compared to HDM-challenged, VC-treated mice, but no difference in *Il17rb* expression was observed between groups (Figure 2D).

**Figure 2.**
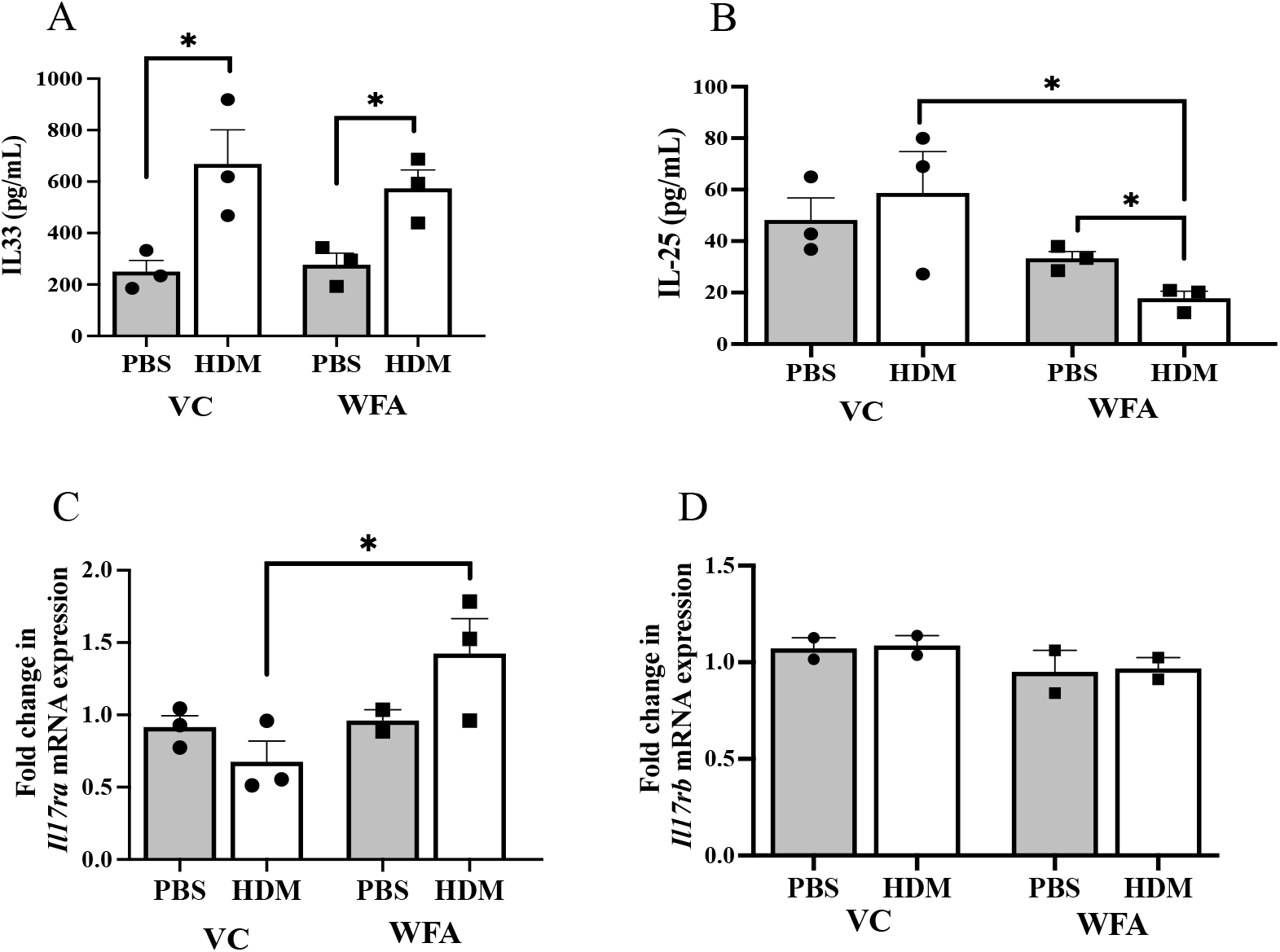
In lung tissue of HDM-challenged mice, WFA reduces IL-25 and increases lung *Il17ra* mRNA expression. (A) IL-33 was measured in whole lung homogenates using ELISA. Data are presented as mean pg/mL IL-33 ± SEM. (n = 3 mice per group). *p<0.050 in HDM-challenged mice compared to saline-challenged mice for both VC and WFA treatment groups. (B) IL-25 was measured in whole lung homogenates using ELISA. Data are presented as mean pg/mL IL-25 ± SEM. *p<0.050 in VC-treated vs WFA-treated HDM-challenged mice (n = 3 mice per group). *p<0.050 in WFA-treated mice comparing saline-challenge to HDM-challenge. (C and D) *Il17ra* and *Il17rb* mRNA expression was measured in whole lung RNA using RT-PCR. Data are presented as fold change in *Il17ra* (C) or *Il17rb* (D) mRNA expression ± SEM relative to *Gapdh* mRNA expression and to the mean of the saline-challenged VC-treated mice. (n = 2-3 mice per group). *p<0.050 *Il17ra* mRNA expression in WFA-treated vs VC-treated HDM-challenged mice.

## Discussion

As an alarmin, IL-25 is pivotal in transducing inflammatory responses to airway epithelial injury. Upon exposure to airborne allergens, airway epithelial cells release IL-25, which induces production of T2 cytokines IL-13, IL-4, IL-5 and IL-9 as well as the eosinophil chemoattractant eotaxin-1 (CCL11). These cytokines and chemokines act to recruit eosinophils and other immune cells to the site of allergen-induced injury (reviewed in Duchesne et al. 2022). IL-25 initiates signaling through a receptor heterodimer comprised of the IL-17RA and IL-17RB receptor subunits (Angkasekwinai et al. 2007). Eosinophils express the IL-25 receptor (Tang et al. 2014); thus, IL-25 can activate eosinophils directly or through T2 cytokine/chemokine production by other airway immune cells. IL-33 is another alarmin secreted by airway epithelial cells following allergen exposure. IL-33 interacts with basophils, natural killer cells, and innate lymphoid cells to induce secretion of T2 cytokines, resulting in recruitment of eosinophils (reviewed in Duchesne et al. 2022). Our data show that, in mice challenged with HDM, WFA reduced airway eosinophilia and inhibited production of lung IL-25 but had no effect on lung IL-33 production. These data suggest a specific effect of WFA on the IL-25 pathway.

Previous studies have demonstrated reduced airway eosinophils with WFA treatment in rodent models of allergic airways disease using ovalbumin (Pretorius et al. 2009; Ali et al. 2023; Zhao et al. 2019). Our study extends the findings of these reports by demonstrating the effect of WFA on airway eosinophilia in a mouse model of acute exposure to the clinically relevant aeroallergen house dust mite, along with reduction of a key T2 cytokine (IL-25) for activating and recruiting eosinophils in human asthma (Tang et al. 2014).

IL-25 signaling occurs through a heterodimer of IL-17RA and IL-17RB subunits, with IL-25 binding directly to IL-17RB, then recruiting IL-17RA to act as a co-receptor. Willis et al. (2015) reported progressive downregulation of lung *Il17Ra* mRNA expression over time in mice challenged with either ovalbumin or cockroach allergen compared to naïve mice. In contrast, we observed that HDM-induced *Il17ra* mRNA expression in lung tissue was increased following WFA administration compared to HDM-challenged mice treated with VC, and we observed no change in *Il17rb* mRNA expression in response to HDM or withaferin treatments. Thus, WFA appears to reverse the effect of allergen exposure on *Il17ra* gene expression in the mouse lung. Further investigation is warranted to determine the effect of WFA on allergen-induced IL-25 pathway signaling downstream of the receptor in lung tissue.

Our study had several limitations, including the small cohort of mice and limited scope of cytokine profiling. Additional work is needed to repeat this study with a larger cohort, as well as to perform lung mechanics measurements of airway response to a methacholine challenge. Future studies will explore the specific effects of WFA on IL-25 signaling in lung cells and the impact of WFA on airway physiology and AHR in the context of allergic airway disease.

## Author contributions

Conceptualization, KA, SP, JLI; conducting of experiments and data acquisition, KA, VLM, JW, MG; data analysis, KA, VLM, JW, JLI; writing and editing of manuscript, KA, VLM, JW, MG, SP, JLI.

## Acknowledgements

The authors wish to thank Haein Kim and Matthew McCravy for their technical assistance in the completion of this study.

## Declaration of Interest Statement

The authors declare that they have no known competing financial interests or personal relationships that could have appeared to influence the work reported in this paper.

## Funding Sources

This work was supported by NIH grants R01HL130234 and R21AI146761 (JLI), India Alliance Wellcome Fund Award IA/CPH/19/1/504607 (SP) and T32ES021432 (MAG, Duke University Program in Environmental Health).

## Notes

### Competing Interest Statement

The authors have declared no competing interest.

